# Reliable Particle Sizing in Vaccine Formulations using Advanced Dynamic Light Scattering

**DOI:** 10.1101/2023.03.22.533677

**Authors:** Coline Bretz, Andrea Jauslin, Dario Leumann, Marius Koch, Andrea Vaccaro

**Affiliations:** LS Instruments AG, Passage du Cardinal 1, 1700 Fribourg, Switzerland; Solvias AG, Large Molecules & ATMP Services, Römerpark 2, 4303 Kaiseraugst, Switzerland

## Abstract

Understanding the impact of lipid nanoparticles’ size on immunogenicity represents an important step for enabling the rapid development of novel vaccines against known or emergent diseases. Dynamic light scattering, also known as quasi-elastic light scattering or photon correlation spectroscopy, has established itself as an optimal analytical method to determine particle size due to its in-situ approach and fast measurements. However, its application to many systems of industrial relevance has been limited due to artifacts arising from multiple scattering. Results interpretation becomes severely compromised depending on the concentration of the system and the size of the particles. In this context, strong sample dilution is often required, bringing additional uncertainties to the formulation development process. Here, we show how advanced dynamic light scattering technology can filter out multiple scattering from the signal and yield fully accurate sizing measurements, regardless of the sample concentration. We illustrate this in a comparative study with standard dynamic light scattering using polystyrene beads as model suspension as well as a concentrated commercial lipid nanoparticle adjuvant (AddaVax™).

## 1. Introduction

Vaccines are among the most remarkable achievements of mankind and continue to be under the spotlight of currently ongoing research [1–3]. Despite the successful eradication of several critical diseases [4] and immense contribution to global public health, there is still a great need for innovative vaccine formulations: common approaches such as the use of aluminum adjuvants or direct injection have proven inadequate for several important pathogens [4]. Challenges inherent to such complex formulations are gradually being overcome as new vaccines continue to appear on the market [5–7], although the scientific developments remaining to be done are considerable [7–10].

The COVID-19 pandemic has brought mRNA technology to the spotlight, and in particular, the use of lipid nanoparticles (LNPs) as a delivery system for mRNA vaccines [8, 11–13]. The advantages of using LNPs for vaccines are numerous, as they are highly customizable, can be manufactured on a large scale, and constitute a safer alternative to viral vectors [8, 14–19]. However, as with any new approach, there remain unknowns and caveats to be exploited. Therefore, accurate characterization is crucial for the complex systems used in human vaccination. Vaccine performance is affected by biophysical properties, and among these properties, LNP size provides a significant contribution to many functional parameters such as biodistribution and cellular uptake; in particular, it determines the entry route of particles into the cells [8, 15]. Typical particle diameters of LNPs are in the order of 10s to 100s of nanometers, can vary widely, and have even been optimized to target certain organs specifically [3, 8, 20–22]. Furthermore, recent studies have evidenced that high antibody titers are generated for an average LNP size of 100 nm in an RNA LNP vaccine in mice [23].

While aluminum salts have long been the only adjuvants included in licensed human vaccines [24–27], engineered LNPs have also emerged as an excellent platform for the delivery of adjuvants [10, 28]. As for their role as nanocarriers, also here the size is a decisive parameter in the adjuvating properties of LNPs. For example, particle size has been shown to impact cell recruitment in a commercial self-emulsifying vaccine adjuvant. Formulations with droplet sizes of 160 nm and 20 nm were compared with a commercial benchmark influenza A adjuvant, MF59. The results showed larger recruitment of immune cells to the injection site for emulsion droplets of 160 nm as compared to smaller particles of 20 and 90 nm size with identical composition [29]. Other adjuvant nanocarriers comprise the AS0 family [4], such as AS03 used in a pandemic influenza vaccine (ca. 150-180 nm emulsion droplets) [22, 29, 30] or AS01 used in Malaria [31] and Varicella Zoster [32] vaccine (ca. 105 nm liposomes) [33, 34].

Knowledge of particle size distribution is not only important for vaccine formulations but also of interest for the characterization of aggregation in biologics, such as for monoclonal antibody (mAb) drugs [35]. While dimer or trimer aggregate formation from mAb monomers has to be strongly controlled, also higher molecular weight (HMW) aggregates can form in the size range of 10s to 100s of nanometers or even micrometers [36]. These issues gain even more importance nowadays as biologics are manufactured at increasing concentrations in order to administer higher doses [35, 37, 38]. With increasing concentration, stability and processibility of the formulations become a major concern due to the higher probability of aggregate formation [39] that is suspected to increase the immunogenicity potential depending on their size [35, 36].

As particle size is one of the key parameters for vaccine efficacy and an important indicator of the quality of biologics, it is crucial to be able to accurately determine the sizes, at best within the original formulation.

A widely used technique for this purpose is Dynamic Light Scattering (DLS)^1^, due to its rapidity, simplicity, low sample volume requirement, high throughput capability, and ability to measure the original formulation in a non-invasive manner (i.e. no column material, no vacuum injection, no dyeing, etc.) [40–43]. Nevertheless, in the case of concentrated systems, DLS can lead to erroneous results as the technique requires highly diluted samples [44–46], a drawback that the majority of analytical techniques have. Standard DLS tools are ill-equipped to provide an accurate size determination in complex, concentrated formulations, such as for vaccines, without substantial sample dilution. However, dilution can be a time-consuming step that may also alter the properties of the sample and therefore lead to erroneous interpretation. In the case of the traditionally used aluminum adjuvants, it has already been shown that dilution severely impacts nanoparticle size [24]. In such cases, reliability in the investigation of native formulations is simply impossible and can substantially impact the quality of the product.

In this work, we detail the challenge of correctly determining the particle size using DLS, especially for sizes in the range of 10s to 100s of nanometers. For this purpose, we experimentally illustrate result differences between a novel technique, named Modulated 3D cross-correlation, and the commonly used DLS approach. The results are obtained from suspensions of polystyrene beads with a diameter of around 132 nm as a model system, as well as the commercial LNP system Addavax™ (approx. 146 nm), similar to currently used vaccine adjuvants [22, 29, 30].

### Dynamic Light Scattering and the Multiple Scattering Limitation

In a DLS experiment, a laser beam is shone onto a sample loaded in a cuvette and the light flux resulting from the interference of the photons scattered by each particle is measured [47]. The measurement principle is schematized in Fig. 1. Due to Brownian motion, the nanoparticles in suspension undergo constant movement. The ensuing continuous rearrangement of the particle positions results in a fluctuation of the measured intensity which allows for the straightforward determination of the diffusion coefficient of the particles under investigation. Small particles diffuse in the medium relatively rapidly, resulting in a rapidly fluctuating intensity signal as compared to the large particles, which diffuse more slowly. Quantitative information regarding the time scale of these fluctuations in the scattered intensity is obtained by the time correlation of the raw signal trace. From there, the diffusion coefficient(s) of the sample is obtained and related to the hydrodynamic diameters of the particles in motion via the Stokes-Einstein equation [47].

**Fig. 1:**
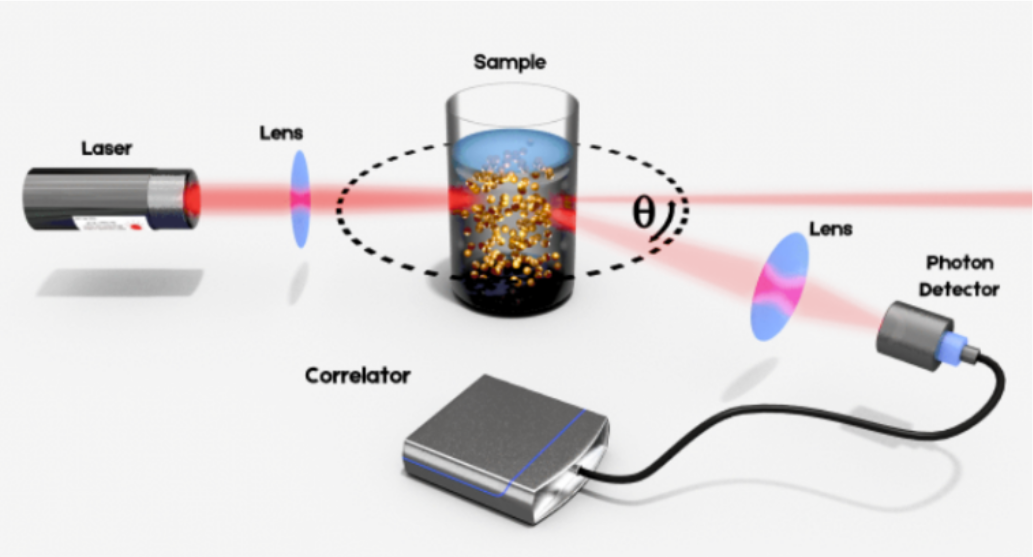
Illustration of the Dynamic Light Scattering principle. The laser beam is focused into the sample suspension containing particles that undergo Brownian motion. The resulting intensity fluctuations in the scattered light are detected at an angle Ѳ for analysis.

This principle, however, requires the measurement of photons scattered only once by the particles: the so-called single scattering events. Apart from highly dilute conditions, multiple scattering may be present in the signal. In such cases, the sizing results will contain significant errors that are undetectable when analyzing a standard DLS measurement. This undetectability is particularly problematic since such an erroneous result can still provide a good signal-to-noise ratio and will thus appear reliable to the analyst.

Mitigation of multiple scattering has been addressed in several DLS instruments through a special “backscattering” detection scheme consisting of measuring the signal scattered at large angles, typically 173°. This scheme allows for a reduction of the path length of the light within the sample, hence reducing the number of multiple scattering events recorded. However, it does not provide a guarantee of avoiding multiple scattering entirely, especially with highly concentrated formulations and larger species due to their increased scattering cross-section. Additionally, one does not know when the sample is dilute enough such that there are no errors present in the scattering signal and therefore additional pre-dilution testing is required.

Novel light scattering techniques developed within the last 15 years have provided a robust solution to this limitation [45, 48]. The general idea is to isolate singly scattered light and suppress undesired contributions from multiple scattering in the experiment. This can be achieved by performing two scattering experiments simultaneously on the same scattering volume, in the so-called 3D cross-correlation scheme. A schematic of the principle of this technique is provided in Fig. 2.

**Fig. 2:**
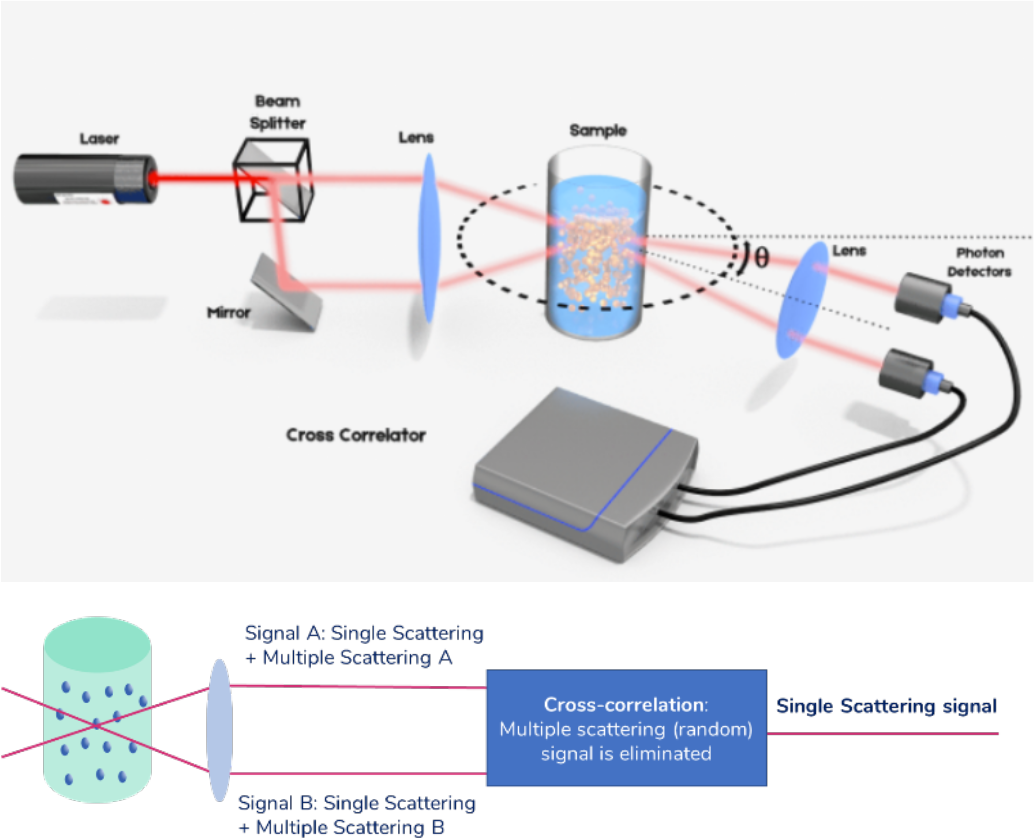
Principle of the 3D cross-correlation technique. Before being focused into the sample solution, the laser beam is split in two parallel beams. The resulting intensity fluctuations in the scattering light are detected similarly as explained in Fig. 1, but using two photon detectors. The analysis is done using a cross correlator. The extension to a scheme in which the illumination beams are alternated and the detection electronics are gated is called Modulated 3D cross-correlation.

In this scheme, the sample is illuminated by two laser beams and the scattered light is harvested by two detectors placed at the same observation angle, above and below the scattering plane. Singly scattered photons follow a deterministic path before being detected, while multiply scattered photons follow a completely random path before reaching the detector. As a result, the two measured multiply scattered photon count signals are temporally uncorrelated. The cross-correlation of the two total count rate signals, within this detection scheme, effectively filters out multiple scattering. This technology offers a fully reliable way of performing DLS experiments, no matter the concentration of the sample. However, the implementation suffers from a reduction in signal quality due to cross-talk between detectors. This scheme has been further improved by temporally separating the two experiments: in the Modulated 3D cross-correlation scheme, [49, 50], the illumination beams are alternately activated with high-speed intensity modulators while the detection electronics are synchronously gated. In this implementation, the quality of the signal is comparable to that of a standard auto-correlation experiment as in standard DLS. This technology effectively removes the upper limit to the concentration of the formulations measured in DLS, allowing for error-free sizing for all kinds of samples, including those showing substantial turbidity. In the following sections, this is illustrated using two types of nanosuspensions.

## 2. Materials & Methods

Measurements in the Modulated 3D cross-correlation scheme were performed using a NanoLab 3D (LS Instruments AG). This device is equipped with a 120 mW laser operating at a 638 nm wavelength. The detection electronics are located at a measurement angle of 90° relative to the incident beam. The particle size was obtained by means of the cumulant fitting method. The results obtained with this instrument are referred to as „Modulated 3D DLS” in the following sections.

To compare the results with those obtained by means of a standard backscattering DLS technology, a Malvern ZetaSizer Nano ZS was used as a benchmarking tool. The device is equipped with a 4 mW laser operating at a 633 nm wavelength. The detection electronics are located at a measurement angle of 173° relative to the incident beam. This backscattering mode minimizes artifacts introduced by multiple scattering. The particle size was obtained by means of the cumulant fitting method. The results obtained with this instrument are referred to as „Standard Backscattering DLS” in the following sections.

12 measurements of 90 seconds were carried out in the Modulated 3D DLS scheme for each data point to maximize results precision (DLS measurements may be as short as 10 to 15 seconds for such samples, which yield a high scattering signal). In the case of the Standard Backscattering DLS, 8 measurements of 10 seconds per repetition were set by the software of the instrument for each data point.

For both instruments, measurement repeatability was within the limits described in the ISO 22412 standard [51].

A volume correction described in [52] was applied to the results. The temperature was set to 25°C in both instruments.

Suspensions of polystyrene beads in water were obtained from Bangs Laboratories Inc. at a concentration of 10.06% (solids) and a nominal diameter of 132 nm.

A squalene-based oil-in-water emulsion of AddaVax™ adjuvant was obtained from InvivoGen, with a nominal particle size of around 146 nm and a concentration of 38.7 mg/mL.

Dilutions were prepared using phosphate-buffered saline (PBS) from Sigma Aldrich. The buffer was prepared by dissolving a PBS tablet in purified water.

## 3. Results & Discussion

### 3.1. Measurements on a model system

In order to conduct experiments on suspensions of polystyrene beads with a nominal diameter of 132 nm, a concentration series between 0.001% and 10% was analyzed. Given the fact that the particles are solid at room temperature and given their high bound surface charge, dilution is expected not to affect the particle size, and hence can be considered a model system.

Fig. 3 shows the sizing results obtained using both DLS measurement methods, together with the error relative to the nominal size.

**Fig. 3:**
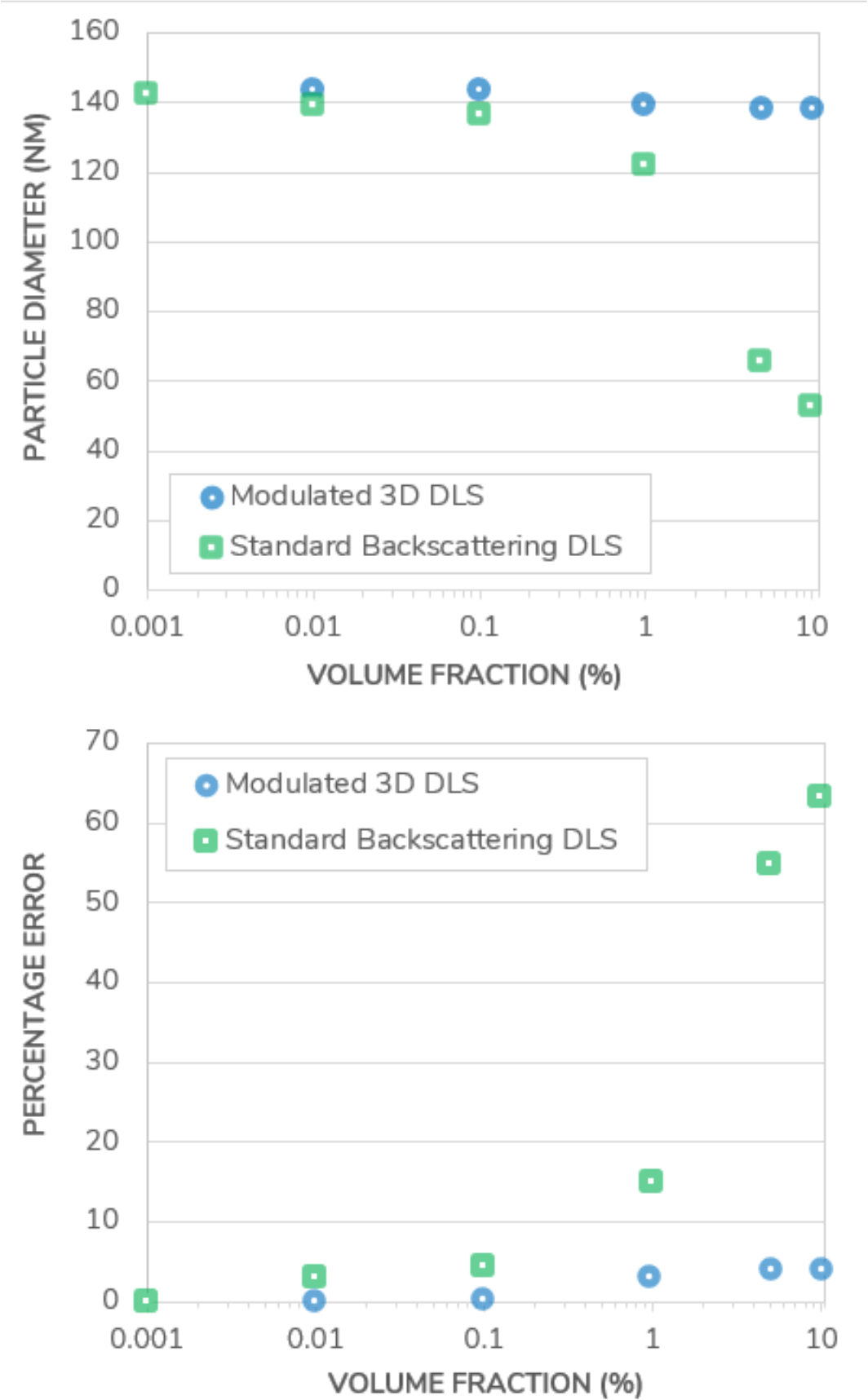
(Top) Particle diameter measured in a polystyrene suspension using Standard Backscattering DLS and Modulated 3D DLS as a function of the volume fraction. (Bottom) Percentage error on the DLS sizing results in both measurement modes, as a function of the volume fraction.

Both instruments produce the same sizing results under highly dilute conditions^2^. Once the concentration increases, the results start to differ substantially: while the measured size does not vary significantly for concentrations lower than 0.1% in the Standard Backscattering DLS scheme, they deviate from the initial size measurement at higher concentrations. This deviation continues to increase with increasing concentration, to reach a situation whereby the measured size is less than half the value of a dilute sample. For example, the measurement error is more than 60% for samples at volume fractions of 10% (Fig. 3 bottom graph). This observed size reduction is typical for the presence of multiple scattering events and, given the attributes of the double layer around the particle thus explaining the difference between the nominal size and the size measured in this work in both DLS instruments in dilute conditions model system under investigation, cannot be ascribed to an actual particle size change. Furthermore, the onset of multiple scattering is already visible at 0.01% volume fraction. Generally, the onset of multiple scattering is difficult, if not impossible, to predict as it requires advanced modeling and a solid knowledge of the optical properties of the suspension. These demands would however put a tremendous barrier to quality control and routine measurements, especially when dealing with multiple complex formulations of differing compositions and properties. As a result, to guarantee trustable measurements when working with Standard Backscattering DLS, there is no reliable alternative to dilution with its accompanying uncertainties of sample alteration.

On the other hand, the size of the polystyrene beads measured in the Modulated 3D DLS mode remains constant, as expected for the polystyrene beads sample under investigation, with less than a 4% variation relative to the nominal value. The Modulated 3D DLS technology thus demonstrates its ability to reliably filter out multiple scattering and to deliver correct and trustable results under any concentration considered.

### 3.2. Measurements on a commercial system: AddaVax™

In the following, a comparison between Standard Backscattering DLS and Modulated 3D DLS is made on a commercial LNP system: AddaVax™, which is a squalene-based oil-in-water emulsion adjuvant with a nominal particle size of around 146 nm, similar to commonly used vaccine adjuvants [22, 29, 30]. The adjuvant was supplied at an initial concentration of 38.7 mg/mL, from which a dilution series was created. A picture of the formulation is displayed in Fig. 4 and shows the turbid appearance of the sample.

**Fig. 4.**
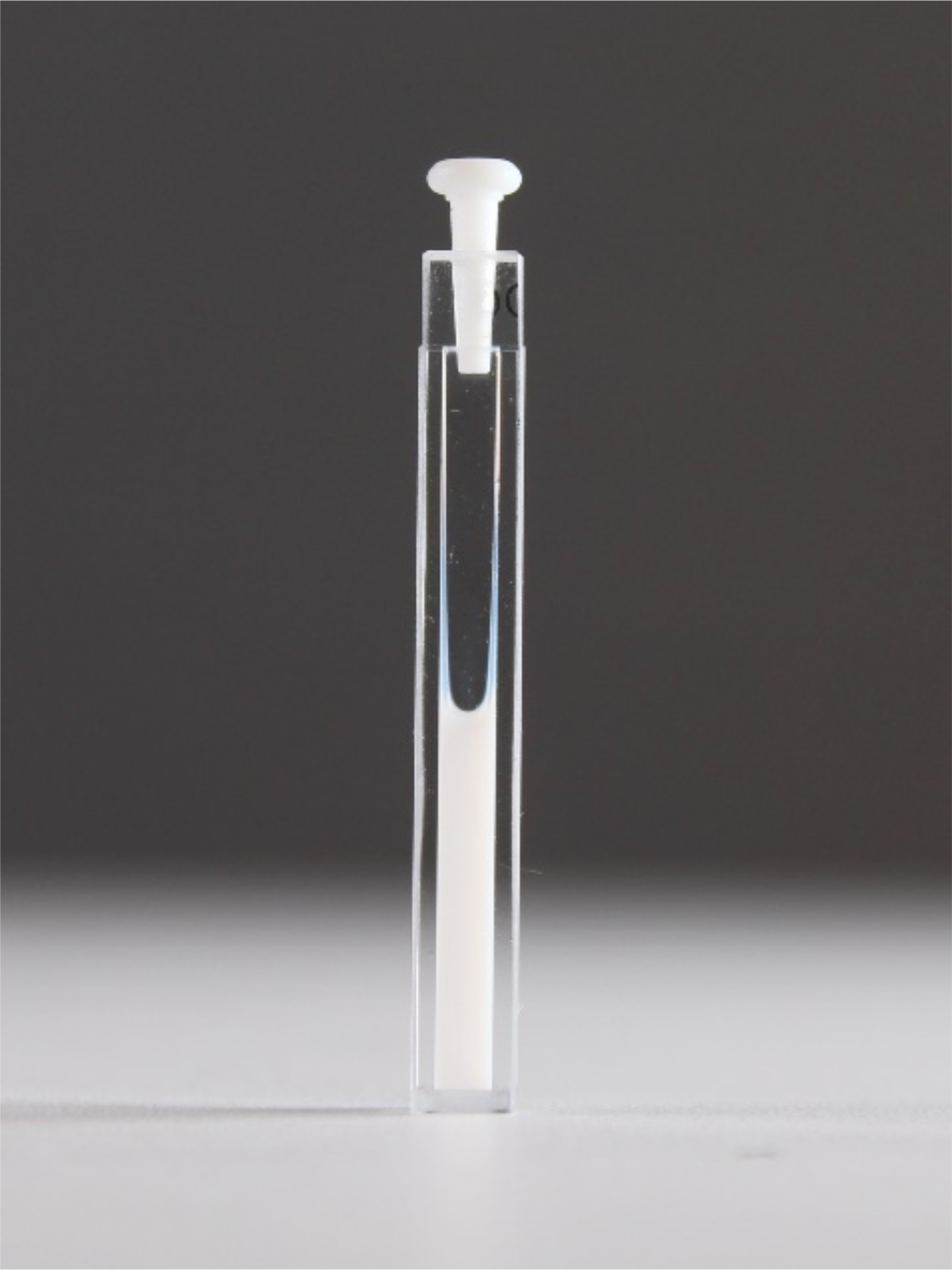
Photograph of AddaVax™ in a measurement cuvette at original formulation concentration (40% volume fraction).

The sizing results, corrected following [52] as in the previous section, are presented in Fig. 5 together with the measurement error relative to the size measured at the lowest concentration.

**Fig. 5.**
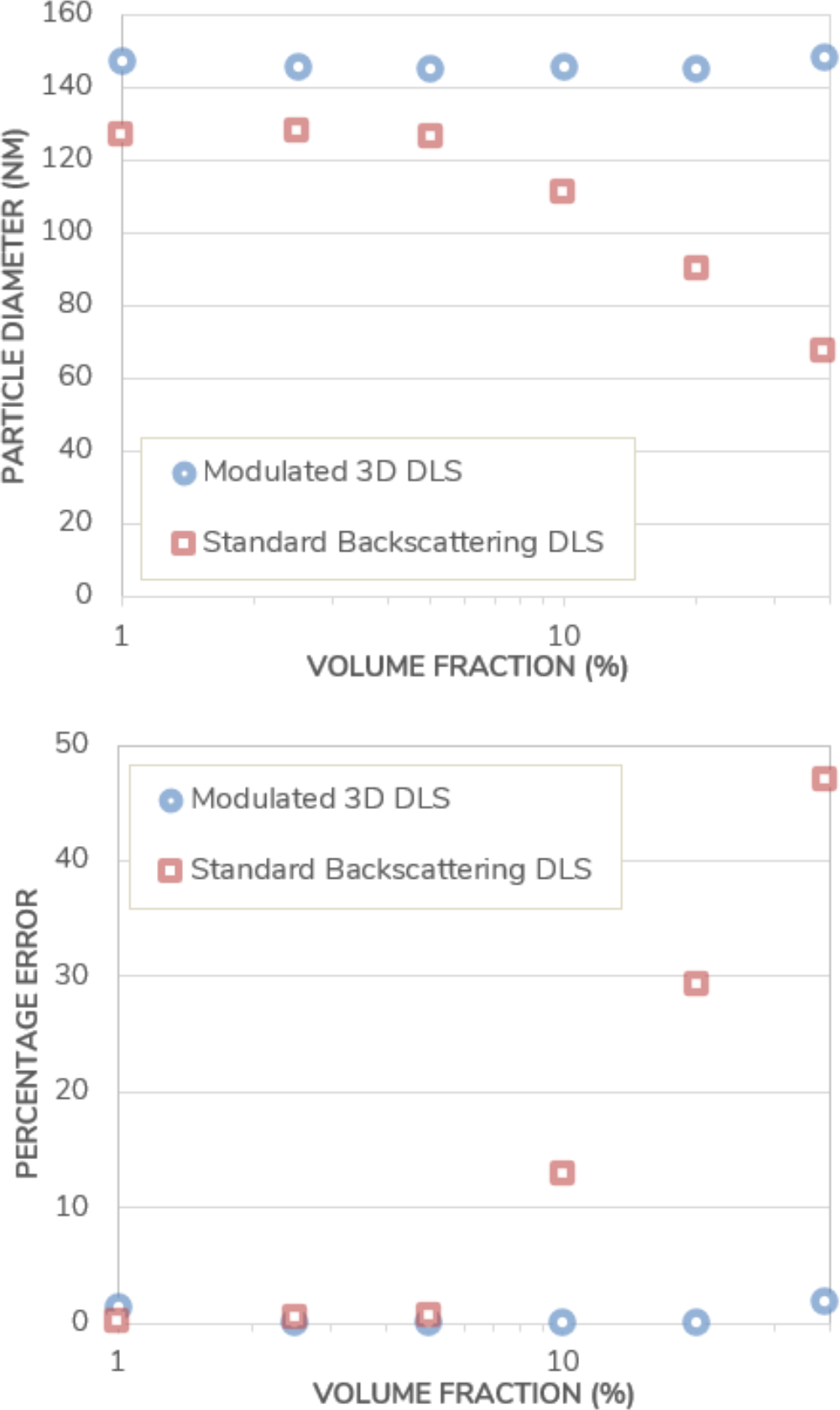
(Top) Particle diameter measured in AddaVax™ using Standard Backscattering DLS and Modulated 3D DLS, as a function of the volume fraction. (Bottom) Percentage error on the DLS sizing results in both instruments, as a function of the volume fraction.

Comparing the results, a size difference of 14% is observed between the two different measuring techniques at the highest dilution. This apparent discrepancy is typical of DLS when applied to polydisperse samples and is not related to multiple-scattering events [53]. In DLS, to obtain the hydrodynamic size, one measures the intensity weighted diffusion coefficient. The active intensity weighting scheme for polydisperse samples changes with the measurement angle. When the measurement angle is decreased, the signal originating from large particles becomes dominant and they contribute more to the result thus leading to an increase in the measured hydrodynamic size. On the contrary, an increase of the angle, as in the backscattering scheme, gives more weight to the smaller particles of the polydisperse sample. The difference is observed in Fig. 5 already for the highest dilution since the Modulated 3D scheme collects intensity at 90° while the Standard Backscattering scheme uses 173°. In comparison, this was not observed in Fig. 3 with a model monodisperse system owing to its very low polydispersity. This emphasizes the importance of the scattering angle chosen, as the Standard Backscattering DLS scheme yields measurement results biased towards lower particle sizes and does not provide an accurate representation of the particle size distribution in the case of polydisperse systems. On the other hand, a measurement angle of 90° provides a more “neutral” characterization of such systems, without bias towards larger or smaller particles.

Similarly, as with the polystyrene beads model system, the measurements results obtained with the Modulated 3D DLS are constant across all concentrations evaluated, indicating that the AddaVax™ suspension is stable against dilution. However, the results obtained with the Standard Backscattering DLS technique show a sharp decrease in the size measured with an onset that begins at a volume fraction just below 10%. As mentioned previously, this onset is not predictable unless resorting to advanced modeling techniques that require precise knowledge of the optical properties of the sample. The measurement error reaches up to 47% in this set of experiments. This illustrates the ability of the Modulated 3D DLS to accurately determine particle size for strongly concentrated samples, while Standard Backscattering DLS fails to do so.

## 4. Conclusion

The field of vaccine formulation is growing and new mRNA technologies are being developed at an ever increasingly fast pace. LNP size is among the most crucial parameters to provide an understanding of improved biodistribution, cellular uptake, and adjuvating potency. In this work, we have shown that standard DLS technology is unable to accurately characterize such formulations. Proper multiple scattering filtering, as introduced with Modulated 3D DLS, is required for fail-safe and trustable measurements in such highly concentrated systems. As an illustration, a polystyrene bead model suspension and a technologically relevant emulsion system (AddaVax™) formulated at high concentration have been investigated. This knowledge similarly translates to high-concentrated biologics formulations with their increased possibility of forming larger aggregates.

With the help of dilution-free measurements of samples, experiment times are significantly shortened and human- or dilution-introduced sample alteration is prevented. The reuse of highly concentrated samples furthermore saves sample material and opens roads for particle size determination of unopened containers as it might be of value for stability studies and quality control of pharmaceuticals. The inevitable presence of multiple scattering events for such a situation, with its erroneous results, can be properly taken care of by using Modulated 3D DLS.

## Acknowledgment

A. Jauslin and M. Koch would like to thank the Solvias AG for the support in carrying out the study.

## Footnotes

1 Depending on the context, DLS is also referred to as quasi-elastic light scattering, QELS, or photon correlation spectroscopy, PCS.

2 The nominal size of 132 nm reported by the particle manufacturer is obtained by means of DLS and by strongly diluting the sample with a 10 mM electrolyte solution. This compresses the electrical

